# From guidelines to practice: Operational criteria for identifying old-growth forests in northern Europe

**DOI:** 10.64898/2026.05.19.724771

**Authors:** Mikko Mönkkönen, Gediminas Brazaitis, Guntis Brūmelis, Bengt-Gunnar Jonsson, Asko Lõhmus, Raisa Mäkipää, Kimmo Syrjänen

## Abstract

Primary and old-growth forests are globally valued for their biodiversity, ecosystem services, and cultural significance. The EU Biodiversity Strategy and EU Forest Strategy for 2030 require strict protection of remaining primary and old-growth forests, yet they cover only about 3% of EU forest area and remain highly threatened. The European Commission’s guidelines define old-growth forests using three main indicators—native tree species, deadwood, and large/old trees—supported by five complementary indicators. Implementing these indicators for boreal and hemiboreal old-growth forests in northern Europe currently lack science-based operational criteria that meet EU legal standards. We provide recommendations for implementing European Commission’s indicators with science-based operational criteria and thresholds to minimize misclassification and ensure cost-effective conservation. Key thresholds include native species dominance, ≥5% deadwood of the total wood volume, and ≥20 large/old trees per hectare. Additional guidance is offered for regeneration patterns, structural complexity, microhabitats, and indicator species, emphasizing that all indicators should be applied collectively.

## Introduction

High conservation value of primary (PF) and old-growth forests (OGF) is recognized in the Kunming–Montreal global biodiversity framework. These forests are globally important for their unique biodiversity, ecosystem services (including carbon storage and sequestration), cultural heritage (especially Indigenous knowledge and rights), and contributions to human wellbeing.

Reflecting this, both the EU Biodiversity Strategy for 2030 and the EU Forest Strategy 2030 commit to defining, locating, and strictly protecting all remaining primary and old-growth forests, regardless of whether Member States have already met their overall 10% strict land-protection target or 20 % target for other conservation areas.

In the EU, PFs and OGFs are rare and threatened, estimated to cover only about 3% of total forest area (Sabatini et al. 2020), and rapidly declining also in the boreal Europe (Ahlström et al. 2022). About 93% of the mapped PFs and OGFs are part of the Natura 2000 Network, and 87% are strictly protected (Barredo et al. 2021). However, due to the mapping deficit we do not know the extent nor the level of protection of the remaining primary and old-growth forests (Sabatini et al. (2020).

“Commission Guidelines for Defining, Mapping, Monitoring, and Strictly Protecting EU Primary and Old-growth Forests” (European Commission 2023) adopts the FAO’s definitions of PF and OGF. PF and OGF are closely related terms, but unlike OGFs, PFs are required to be large enough to maintain natural ecological processes and may include younger natural successional stages that result from those processes. According to Commission guidelines an OGF “is a forest stand or area consisting of native tree species that have developed, predominantly through natural processes, structures and dynamics normally associated with late-seral developmental phases in primary or undisturbed forests of the same type.” The Commission guidelines explicitly allow historical management when it has not significantly altered current structures but excludes stands that “are under active productive management”. Thus, a major distinction from PFs is that an OGF can be a stand of any size that have late-seral structures (legacies from the past) present, whereas natural processes may not all be currently active there. Our focus here is to identify the sets of such structures to enable effective detection of OGF based on scientifically defendable operational criteria (detection rules).

Precise identification rules for PF and OGF are also required across EU regulations. The Renewable Energy Directive (RED III) prohibits harvesting of PF and OGF and requires strict protection of these. The EU Nature Restoration Law does not mention PF or OGF explicitly but forbids altering Habitat Directive forest types in good condition. The OGF criteria presented here may help assess such conditions.

There is clearly a continuum between OGF and non-OGF with a grey area in between that could be considered as future OGF. To govern this ambiguity, the Commission has developed a list of indicators for OGFs (Table 1) that the Member States must cling to while creating methodologies and operational criteria to guide OGF delineation in practice (European Commission 2023). The Commission necessitates that the methodologies are transparent, publicly available, and science-based, enabling cross-border consistency and objective verification by stakeholders (European Commission 2023). While the identification of OGFs is grounded in ecological science, its application is inherently a governance process. Even though OGFs cover only a small proportion of productive forest area in the EU, criteria, thresholds, and interpretations mediate conflicts between conservation obligations, forest ownership, industrial use, and rural livelihoods. Thus, developing the operational criteria is not merely technical but institutional and political in its consequences.

**Table 1.** EU Commission’s indicators and their description for identifying old-growth forests (OGF), as well as our recommendations how to apply them in European boreal and hemiboreal forests. Forest stands qualify as OGF if all three main and at least two complementary indicators are met.

|  | European Commission's description | Recommendations for European boreal and hemiboreal forests |
| --- | --- | --- |
| **Main indicators** |  |  |
| 1. Native tree species | *OGF are composed of native species. However, the presence of a small number of non-native trees should not disqualify a forest from being designated as old-growth, if they do not significantly disturb ecological processes.* | Only stands dominated by native tree species should be included for further consideration. |
| 2. Deadwood | *OGF are characterised by a high proportion and diversity of standing and lying deadwood. The amount and type of deadwood can vary greatly between OGF (depending on the forest type, the local environmental conditions, and the area's recent disturbance history).* | Any stand where the proportion of deadwood, with >10 cm diameter, out of the total tree volume (alive and dead trees) is above 5 % should be considered a potential OGF. Moreover, for a stand to be considered OGF, deadwood diversity must be present with standing dead wood and more than one decay stage, including well-decayed deadwood represented. |
| 3. Old or large trees | *OGF are often characterised by a high volume of standing trees relative to earlier development stages for the given forest type and local growing conditions, and by the presence of old or large trees, some of which may reach the maximum age known for the species under the local site conditions.* | Any forest stand where there are more than 20 trees/ha older than indicated in Table 4 for different regions and site types should be considered a potential OGF. |
| **Complementary indicators** |  |  |
| 4. Natural regeneration | *Most OGF stands originate from natural regeneration, but some sown or planted forests can meet the definition, if given enough time to develop the characteristics of OGF.* | Include all naturally regenerated stands, but planted or sown stands only in exceptional cases. |
| 5. Structural complexity | *OGF are generally characterised by structural complexity. This can include a multi-layer canopy structure, horizontal structural diversity, and soil micro relief structures such as mounds caused by uprooting.* | Include stands with horizontal (natural regeneration gaps) and vertical diversity in the tree layer. If an old tree cohort is dominant, trees significantly younger are reaching up to the canopy; if a younger tree cohort dominates, the presence of old trees suffices (main criterion 3). |
| 6. Habitat trees | *OGF are often characterised by the high density and high diversity of tree-related microhabitats. These are defined as a distinct, well-delineated structure occurring on living or standing dead trees, that constitutes a particular and essential substrate or life site for species or species communities during at least a part of their life cycle to develop, feed, shelter or breed.* | Include stands with visible occurrence of tree-related microhabitats. Most important are: tree/woodpecker holes, hollow trees, kelo trees, fire scars. |
| 7. Indicator species | *OGF often host species of late-seral developmental phases that are specific to a certain forest type. These can include* | The presence of species that is reliably demonstrated to be OGF specific |
|  | *species on the red-list of the International Union for Conservation of Nature(IUCN).* | indicates that the forest stand possesses OGF qualities. We recommend (i) using pre-defined national species lists containing at least, and preferably more than, 20 such species that occur in multiple types of OGF across the region and, collectively, represent a range of OGF conditions and taxon groups, and (ii) reducing random errors by requiring that reliable records of at least two listed species within the last 20 years must be present for a stand to qualify as OGF under this criterion. The absence of the listed species must not be interpreted as a non-OGF stand. |

Detecting OGFs as stands with specific structures in forest landscapes requires context-dependent interpretation of Europe’s diverse environments, forest dynamics, and histories of use. Boreal and hemiboreal forests in northern Europe merit separate scrutiny from temperate forests because their natural dynamics and resulting structural features are distinct and well documented. Science-based operational criteria are essential for navigating political interests in this region. This article offers the first recommendation for regional harmonisation of OGF detection rules across northern Europe, including Estonia, Latvia, Lithuania, Sweden, Finland, and Norway, still acknowledging the variation among biogeographic zones and forest types in the region.

The aim of this article is to establish science-based operational criteria for OGFs in boreal and hemiboreal region to support their mapping and protection. We first review the natural disturbance regimes that, together with regional species pools and historical factors, have shaped - and continue to shape - primary and old-growth conditions and variation therein. We then outline conceptual and methodological challenges in defining operational OGF criteria and propose solutions. Following EU guidelines, we present concrete operational criteria based on evidence from remaining old-growth forests and their natural dynamics. Finally, we provide insights for the implementation of the criteria, highlight remaining knowledge gaps and ways to address them, and discuss governance, institutional, and social aspects of OGF detection and protection.

### Dynamics and structure of the European boreal forests

Although much of Europe has been shaped by millennia of land use, northern European forests still retain landscapes with relatively limited human impact (Kuuluvainen et al. 2017). These legacies, particularly in Fennoscandia, have been extensively studied (Esseen et al. 1997), providing a strong understanding of how natural processes shaped forest structure and biodiversity (e.g. Berglund & Kuuluvainen 2021). In the boreal–temperate transition zone (hemiboreal zone), natural dynamics were modified earlier, yet OGF features and biota remain (Nilsson 1997; Lõhmus & Kraut 2010). Thus, despite today’s substantial human influence, documented baselines exist for defining operational OGF criteria.

Following the last glacial period (∼18,000–10,000 years ago), species gradually recolonized northern Europe, making boreal and hemiboreal species assemblages relatively “young” and dynamic (Mönkkönen et al. 2018). Glacial history also explains the comparatively low diversity of trees (Huntley 1993), birds, and mammals (Mönkkönen & Viro 1997) relative to other boreal regions in the eastern Palaeartic or Neartic. Nevertheless, the region hosts tens of thousands of often highly specialized multicellular species forming distinct ecosystems.

The present distribution of European boreal forests developed during the Holocene. Ahti et al. (1968) distinguished four zones (Fig. 1) based on mesic-site plant communities, showing increasing conifer dominance and declining temperate broad-leaved trees from south to north.

**Figure 1.**
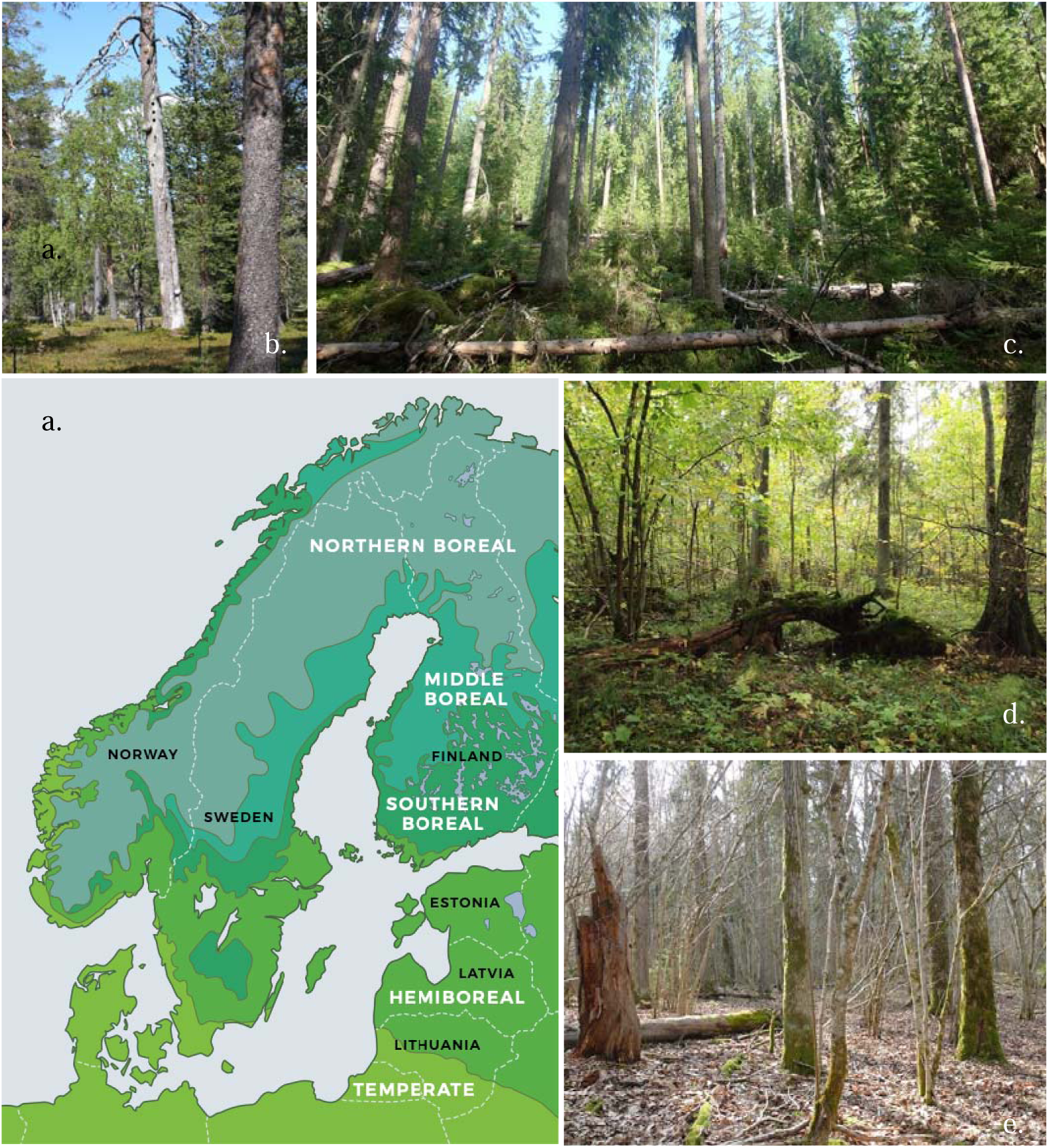
Panel a: Map of the boreal forest zones in northern Europe. Redrawn form Ahti et al. 1968. Examples of old-growth forests from different zones: Panel b: northern boreal pine-dominated forest in Pellokielas, northern Sweden, including a standing pine snag, aka kelo tree. Photo: B.G. Jonsson. Panel c: southern boreal spruce-dominated forest in Evo strict forest reserve, southern Finland, with multilayered structure and a small gap with regenerating spruces. Photo: G. Brazaitis. Panel d: Aegopodium-type herb-rich mixed forest, showing gap-phase dynamics, multi-layered canopies, and large decayed fallen trunks. Järvselja Nature Reserve, East Estonia. Photo by A. Lõhmus Panel e. Broadleaved (mixed-oak) hemiboreal old-growth in western Latvia with spruce understorey. Photo: Sandra Ikauniece.

Boreal forests are influenced by a wide range of disturbances varying in size, severity and frequency, including fire, wind, snow damage, flooding, insects, fungi, and browsing (Esseen et al. 1997). Earlier emphasis on frequent, high-severity, stand-replacing disturbances has shifted, and evidence shows that natural disturbance regimes are more heterogeneous and dominated by low-and moderate-severity disturbances (Kuuluvainen 2009; Kuuluvainen & Aakala 2011; Kuuluvainen et al. 2015). Key disturbance types, in decreasing prevalence, include (Berglund & Kuuluvainen 2021):

- small-scale gap dynamics
- cohort dynamics from partial disturbances such as surface fires and windthrow
- mixed-severity fire regimes
- stand-replacing disturbances

Even large fires are less uniform and less fully stand-replacing than earlier believed (Kneeshaw et al. 2011). Similar regimes operate in Russian boreal forests (Shorohova et al. 2009; Khakimulina et al. 2016) and become even more dominated by small-scale gap dynamics toward the Baltic region because forest fires are less common in the hemiboreal than further north (Acil et al. 2025, Manton et al. 2025).

Current OGFs represent a legacy of these processes: heterogeneous structure, old trees, and specialized species depending on long-term continuity (Nordén et al. 2014). Because gap dynamics and partial disturbances dominate, natural landscapes would contain a high proportion (≥50%, often 70–95%) of stands with old trees (Berglund & Kuuluvainen 2021). In these landscapes, stands are typically uneven-aged, multilayered, and structurally complex, with reverse-J age distributions where younger trees dominate. Abundant deadwood is typical for natural boreal forests. After intense disturbances deadwood volumes might exceed hundreds of m^3^·per hectares. After these pulses, deadwood amounts decline over 50-100 years before gradually increasing (Löfroth et al. 2023). Considerable structural variation exists among vegetation zones, productivity classes, and dominant tree species (Skarpaas & Halvorsen 2022; Syrjänen et al. 2024), requiring careful design of operational criteria for OGFs to avoid misclassification.

### Theoretical background for defining criteria

The methodological challenge of detecting OGF in the spirit of the EU Biodiversity Strategy involves three overarching considerations. Together, they mean that we need to develop ways to *interpret* measured forest characteristics to include OGFs as completely as possible.

First, detecting “all remaining” OGF requires accounting not only for typical stands, but for the *full variation* of both OGF and non-OGF across environmental gradients, to achieve reliable discrimination. Second, the European Commission (2023) restricts OGF detection rules to seven *pre-defined indicators*, of which only three are mandatory and two (structural complexity and indicator species) allow particularly broad interpretation in qualitative terms (Table 1). Furthermore, two formerly used boreal OGF indicators (e.g., Rouvinen & Kouki 2008; Martin et al. 2023) are explicitly relaxed, permitting limited past management and including secondary stands. Third, mapping of OGF stands is mainly based on field surveys where the indicator values are measured using a variety of field sampling methods. These data are always influenced by variation due to *measurement errors* stemming from random factors (e.g., exact measurement location), sampling design (e.g., number and size of sampling units) and human errors.

Detection theory combined with decision theory (Hautus et al. 2021) provides useful tools for formalising trade-offs and uncertainties inherent in using simplified indicators. Such indicators inevitably generate both *false negatives* (missed OGF, with ecological costs) and *false positives* (non-OGF incorrectly classified as OGF, with economic or opportunity costs; Fig. 2). The magnitude of these costs depends not only on indicator accuracy but also on the relative abundance of non-OGFs and their structural similarity to OGF. For example, even if indicators correctly classify 90% of OGF and 99.1% of non-OGF, a situation in which non-OGF constitute 99% of the landscape would result in half of all selected sites being non-OGF, while 10% of true OGF would be missed. Threshold setting thus involves two interacting *trade-offs*: between ecologically versus economically costly decision errors, and between survey costs and classification accuracy, as comprehensive surveys reduce uncertainty but require greater effort.

**Figure 2.**
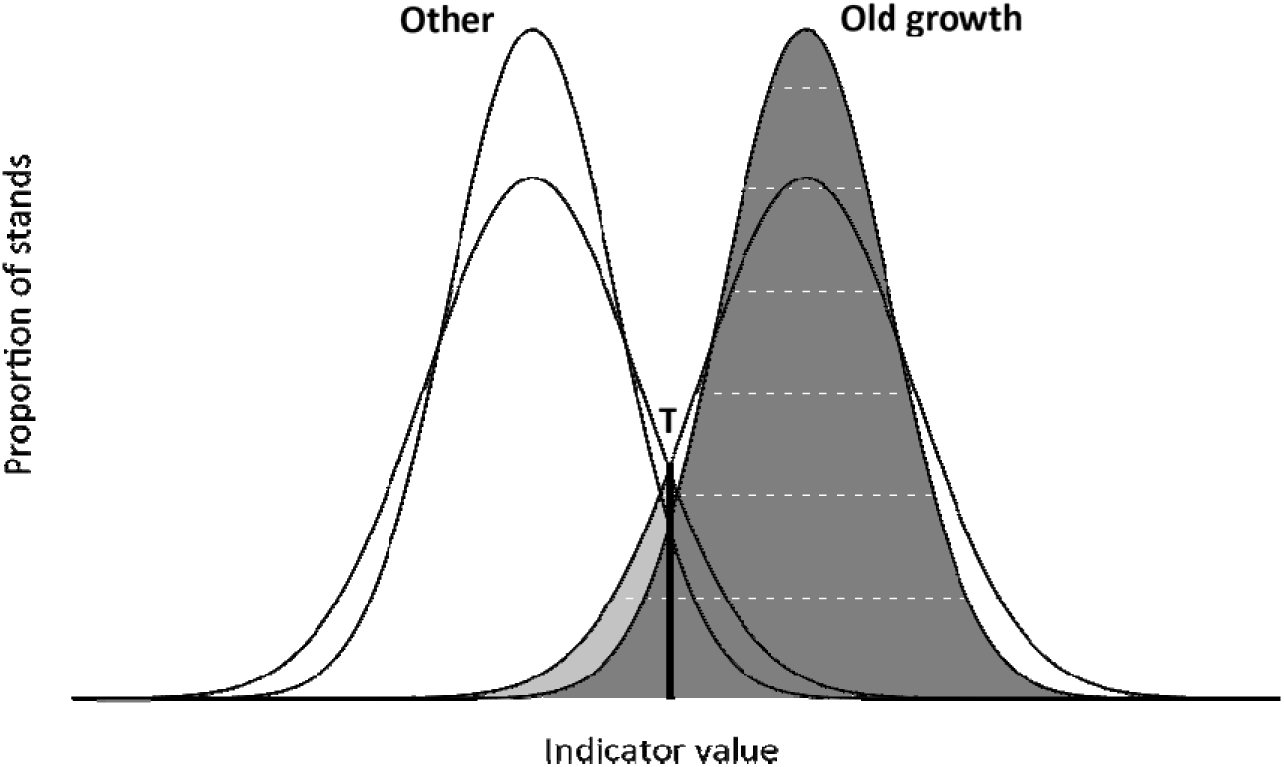
Basic detection problem of OGF (dark grey distribution on the right) from ‘other’ forests (left) based on a hypothetical normally distributed indicator with partial discrimination (here 10% shared area between distributions). Setting a threshold (T) midway between the means excludes 5% of true OGF stands (false negatives) and includes 5% of other forests (false positives). However, if the actual indicator measurements are with an error, both the variance of the distributions, the overlap, and the share of classification errors are expected to increase (lower-peaked distributions; additional false negatives shown in light grey). Shifting the threshold to reduce one type of classification error inevitably increases the other, whereas the absolute number of false detections depends on the error rates and the total number of stands.

The European Commission (2023) guidelines imply a decision rule that *combines multiple stand-level criteria,* a theoretically sound approach for reducing uncertainty. For example, three fully independent indicators with the characteristics used above would almost eliminate false positive detections when applied as mandatory indicator, potentially allowing lower thresholds that miss fewer true OGF. However, combining multiple mandatory indicators increases the risk of false negative errors, as failure to meet any one threshold leads to exclusion. In contrast, complementary (OR-type) indicators reduce the risk of missed detections, as a stand is identified if it meets at least one threshold, reflecting an ecologically sound idea that OGFs may differ in their value sets. This corresponds to the approach to combine mandatory (main) and complementary indicators.

Consider again a landscape where OGF constitutes 1% of all forest and suppose we aim for 99% of final detections to be true OGF to minimise economic costs. Where exactly this condition is met depends on actual distributions and overlaps in indicator values and can be solved numerically.

Thus, a single hypothetical normally distributed indicator with means of 100±25 (SD) and 20±10 in OGF and non-OGF forests, respectively (loosely reflecting typical deadwood volumes ha^-1^), yields a threshold at 57, where 4.5% of true OGF are missed. However, increasing the standard deviation by a factor of 1.5 (to 37.5 and 15, respectively) shifts the threshold value to 77 and increases the false negative rate to roughly 27%. Using three fully independent and mandatory indicators with identical distributions (for simplicity) would allow the threshold at about 37 that misses only 2.4 % of true OGFs. With the increased variability, the corresponding threshold increases to about 46, with approximately 20.6% of true OGFs missed. In contrast, complementary (OR-type) use of multiple indicators would improve detection by reducing the probability that all indicators simultaneously miss the same stand, but it requires stricter thresholds to maintain the 99% positive predictive value condition. Thus, precise measurement, low variation and proper use as mandatory or complementary indicators are important indicator qualities.

Omission rates would rise considerably if indicator independence assumption was relaxed. Notably, several OGF indicators share common components related to forest age and productivity in the boreal zone (e.g. Hämäläinen et al. 2024). However, the main indicators (presence of native tree species, tree age and deadwood amounts) are unlikely to be strongly correlated in old forests. For example, Skarpaas & Halvorsen’s (2022) extensive literature review showed total deadwood volume is more strongly related to site productivity, dominant tree species, and past management than to mean tree age. We therefore conclude that using a combination of the three mandatory indicators and a set of complementary indicators reduces omission rates to an acceptable level.

Verifying this assumption and calculating precise false negative versus false positive error rates would, however, require comprehensive data from both verified OGFs and mature forests representing different biogeographical zones, site types, and management histories, which is typically missing.

For these reasons, our approach to the OGF operational criteria and threshold values was guided by four qualitative assessments.

i. *Presence of sufficient baseline data* to understand full OGF variation across climatic zones and forest types. While specific features (e.g. fractions of live trees or deadwood) may have emerged as diagnostic in case studies, extensive regional documentation is needed to derive broadly applicable OGF-specific thresholds.
ii. *Methodological robustness* in quantifying indicator values and their confounding factors (such as site productivity). Notably, if the presence of certain structures or features is sufficient, this may be more robust than quantitative measurement in variable surveys.
iii. *Omitting the structurally poorest decile of OGF measurements* when considering quantitative thresholds. This addresses the risk that atypical values reflect measurement error in noisy datasets (e.g. small plots) or relaxed criteria in initial OGF assignments (“the best found” for a particular region; see Lõhmus and Kraut 2010). Given the EU requirement to protect all OGFs, we avoided economic arguments to reduce false positives and considered that missing the structurally poorest 10% of true OGFs may entail a relatively low ecological cost.
iv. *Presence-only significance* of OGF-specific species (if based on recent, reliable, and accurately located record) reflects multiple important ecosystem qualities. In contrast, absence of such species is not considered, as it may result from either unsuitable habitat, population fluctuations, or non-detection.

### Operational criteria to identify the remaining OGFs in the boreal and hemiboreal Europe

This section reviews European Commission’s indicators for identifying old-growth forests. Forest stands qualify as OGF if they meet all three main indicators plus at least two complementary ones (Table 1). Such combinatory use of indicators - as discussed above - effectively reduces the risk for both false negative and false positive classification of stands. Table 1 provides our recommendations for applying these indicators in the boreal and hemiboreal Europe.

The Commission guidelines explicitly allow historical management. Past drainage is common in the region (Paavilainen & Päivänen, 1995), and although it alters species composition, such forests may regain structures and species typical of OGFs (Remm et al. 2013). Therefore, previously drained forests can still be considered potential OGFs.

### Main indicators

#### 1. Native species

Native tree species are those that colonized the region without human assistance after the last glaciation. Their presence reflects long biogeographical continuity and local adaptation (Buchwald 2005; Svenning & Skov 2007). In boreal and hemiboreal Europe, introduced species form only a minor share of the growing stock and are mostly confined to experimental stands (e.g. Sander & Meikar 2009), apart from *Pinus contorta* in Sweden, which covers ∼2% of productive forest land (SLU 2024). We recommend including only stands *dominated* by native tree species for further consideration (Table 1).

OGFs in this region are dominated by long-lived native species adapted to natural disturbance regimes and regional conditions. Across countries, 24–30 native tree species occur. Scots pine and Norway spruce dominate, together accounting for 70–80% of the growing stock in Fennoscandia and 50–65% in the Baltic states (Table S1). Birches contribute about 16%, while other broadleaved species form small volumes but are important for biodiversity due to their unique associated biota.

#### 2. Deadwood

Natural disturbances typically create and maintain large amounts and a high diversity of deadwood. Absolute deadwood volumes among OGF stands vary greatly - on average from 20 to 150 m3/ha (Table S2) - with vegetation zone, site productivity and dominant tree species, reflecting differences in disturbance history, growth, mortality and living tree volume (Siitonen 2001; Skarpaas & Halvorsen 2022; Syrjänen et al. 2024).

Because of this, quantitative thresholds based on absolute deadwood volume are difficult to define, and high thresholds risk false negative classifications, while low thresholds increase false positive ones. In contrast, the *relative proportion* of deadwood of total tree volume (dead and living) varies much less, making it a more robust indicator for OGFs (Skarpaas & Halvorsen 2022; Syrjänen et al. 2024). Some differences exist between spruce and deciduous vs pine forests, but not strongly among boreal subzones (Table 2).

**Table 2.** Mean, standard deviation and 10th percentile (decile) of dead wood proportion (%) of total wood volume (living and dead) in old growth deciduous-, spruce- and pine-dominated (pine bogs not included) forests in three large compilations of OGFs (n = number of stands; * recalculated from original data) representing biogeographic zones of boreal forests.

| Boreal zones | n | mean | std | 1<sup>st</sup> decile | Source |
| --- | --- | --- | --- | --- | --- |
| Deciduous dominated |  |  |  |  |  |
| Hemiboreal | 21 | 26 | 14 | 7 | Rosenvald et al. 2026* |
| Southern and middle boreal | 22 | 16 | 8 | 2 | Siitonen et al. 2026 |
| Spruce dominated |  |  |  |  |  |
| Hemiboreal | 33 | 34 | 12 | 17 | Rosenvald et al. 2026* |
| Hemi- and southern boreal | 24 | 25 | 13 | 11 | Syrjänen et al. 2024 |
| Middle boreal | 25 | 31 | 10 | 19 |  |
| Northern boreal | 39 | 28 | 13 | 15 |  |
| Southern and middle boreal | 121 | 15 | 9 | 4 | Siitonen et al. 2026 |
| Pine dominated |  |  |  |  |  |
| Hemiboreal | 19 | 18 | 13 | 5 | Rosenvald et al. 2026* |
| Hemi- and southern boreal | 10 | 10 | 5 | 5 | Syrjänen et al 2024 |
| Middle boreal | 10 | 35 | 10 | 19 |  |
| Northern boreal | 22 | 18 | 7 | 10 |  |
| Southern and middle boreal | 90 | 16 | 15 | 4 | Siitonen et al. 2026 |

The first decile of relative deadwood volume in OGFs ranges from 2 to 19 %, and in most cases the value is larger than or equal to 5% (Table 2). To avoid excessive false negatives and to get closer to European Commission’s objective to protect all remaining OGFs, we recommend considering potential OGFs all forests with relative deadwood volume greater than 5 %.

However, it is important to bear in mind that some forests can be OGFs even if below the 5 % deadwood threshold as Table 2 shows. Examples include pine dominated southern and hemiboreal forests, which are frequently shaped by historical selective cuttings (Lõhmus & Kraut 2010).

Likewise, Syrjänen et al. (2024) reported low relative deadwood volumes in many northern boreal OGFs. In both cases, these forests should still be considered OGFs according to the Commission’s definition (see above).

In similar vein, many mature stands are not OGFs even if the relative deadwood volume is above 5 %. In managed mature forests with recurring thinnings, total tree volume typically remains lower than in OGFs, and thus, 5 % threshold can be met with lower absolute deadwood volume. These stands can be disqualified based on other indicators. We therefore urge to use the deadwood indicator flexibly and always in combination with other indicators, i.e. including some forests below the threshold into OGFs and excluding others above the threshold when the other indicators so imply.

In addition, it is necessary to consider deadwood quality (Table 1). Deadwood diversity is a typical OGF characteristic: roughly one-third standing, two-thirds downed; about one-third large dimension; and about one-third heavily decayed (Shorohova & Kapitsa 2015; Skarpaas & Halvorsen 2022). Although variable, the presence of diverse deadwood - both recent and well-decayed material and standing and downed - serves as a useful complementary qualitative indicator.

#### 3. Old or large trees

A key indicator of OGFs is the presence of old trees, not the dominance of old cohorts. Natural disturbance regimes—especially cohort dynamics and mixed-severity fire—commonly produce OGFs in which younger cohorts dominate the canopy (Nilsson et al. 2002; Lõhmus & Kraut 2010). Thus, the defining feature is the persistence of an old legacy cohort. Therefore, using the age of the dominant cohort as a decisive criterion for identifying OGFs is inappropriate and contradicts the European Commission’s definition, which explicitly includes stands exhibiting visible signs of abiotic (storms, snow, drought, fire) and biotic (insects, disease) damage. In European boreal forests, densities of at least 20 old living trees per hectare have been reported (Nilsson et al. 2002). Thus, we recommend that any forest stand where the density is higher than 20 trees/ha older than indicated in Table 3 for different regions and site types should be considered a potential OGF.

**Table 3.** Minimum age limits (in years) for trees considered old in different forest habitat types on mineral soil and biogeographic zones of boreal forests. The same age limits are applicable for forests on peat soils.

|  | Xeric | Subxeric | Mesic and herb-rich |  |
| --- | --- | --- | --- | --- |
|  |  |  | Conifer | Deciduous |
| North boreal | 180 | 160 | 140 | 80 |
| Middle boreal | 160 | 140 | 120 | 80 |
| Southern and hemiboreal | 140 | 120 | 120 | Aspen, birch & black alder 80<br>Other species 120 |

There is no single threshold after which a tree becomes old. We adopted here a commonly used age limit to consider a tree “old” when the age is more than 1.5 times typical species-specific silvicultural rotation age (Kouki et al. 2018). We used recommended rotation ages (Äijälä 2019), multiplied them with 1.5 and rounded to the nearest 20 years. The selected practical age limit has an biological reasoning as the silvicultural rotation age reflects physiological growth processes and is determined by biological culmination point (Assmann 1970). This age limit varies depending on the type of site, geographical location and dominant tree species (Table 3).

Although the Commission notes that OGFs may exhibit high standing volume (m^3^/ha), this is often not the case in boreal and hemiboreal regions, because remaining OGF stands frequently occur at higher elevations or on poor soils (Virkkala & Rajasärkkä 2007; Angelstam et al. 2020), and in OGFs on productive soils, a significant proportion of wood volume can reside in dead trunks (Rosenvald et al. 2026). Stand volume is therefore an unreliable indicator in boreal Europe due to their strong dependence on site productivity.

### Complementary indicators

#### 4. Natural regeneration

Natural regeneration is a key complementary indicator because it reflects the long-term continuity of locally adapted tree populations. In naturally regenerated OGFs, stand structures are shaped by disturbances and successional processes, producing features such as large, old trees, diverse deadwood, and high structural complexity (Peterken 1996; Kuuluvainen &Aakala 2011). In contrast, artificially regenerated forests typically exhibit simplified structures and slower development of OGF attributes, even when composed of native species. (Buchvald 2005; Kuuluvainen 2009).

Nevertheless, some artificially regenerated stands can qualify as OGFs. For example, coastal pine stands in Latvia — like those planted in the 1700s to stabilize dune systems (Sarma 1974) —have developed OGF characteristics over centuries (Brumelis et al. 2005). Comparable cases exist in Sweden, where old coastal pine plantations occur on sand dunes. Although often influenced by historical management and limited in extent, such stands may be classified as OGFs when main indicators are met, and recent human intervention is absent. Our recommendation is to include all naturally regenerated stands, but planted or sown stands only in exceptional cases.

#### 5. Structural complexity

Horizontal and vertical heterogeneity is limited in managed forests where the aim is to produce a homogenous stand of evenly spaced trees of similar size. By contrast, natural disturbances such as forest fire, windthrow and mortality due to insects and fungi create significant structural variation typical for OGF. According to the natural disturbance regimes outlined above, structural heterogeneity is closely linked to their impact on the tree layer. Gap dynamics open small gaps allowing new trees to establish and by filling gaps create a multilayered canopy. Low-severity surface fires leave a fraction of older trees alive while allowing regeneration of a new cohort of trees and on mesic sites support regeneration of pioneer deciduous species. Larger mixed-severity fires may kill a significant fraction of trees locally but often leaving surviving patches of trees at a larger scale. Hence and regardless of the disturbance factor, this creates stands with significant horizontal and vertical variation, as well as uprooted trees, typical for old-growth conditions (Fig. 1).

#### 6. Habitat trees

An additional indicator of old-growth forest is living and dead habitat trees supporting high diversity of microhabitats, which are considered to provide a “substrate or life site for species or species communities during at least a part of their life cycle” (Larrieu et al. 2022). Many studies confirm the association of these microhabitats with diversity and abundance of forest species, with the most studied being on tree cavities due to direct association with birds (Nirhamo & Kouki 2025). Other well-known associations include crown dead wood with saproxylic beetles (Gossner et al. 2013), and new research is adding knowledge on the importance of a wide range of microhabitats for micro-invertebrates (Majdi et al. 2025). Barkless standing dead Scots pines, aka kelo trees, are a long-lasting, ecologically integral part of the natural boreal forest supporting a variety of species. Because of their longevity and extremely slow turnover dynamics and importance for biodiversity, protection of the remaining kelo trees should be of high priority (Kuuluvainen et al. 2017).

The relationship between tree-related microhabitats has not been extensively studied in old-growth forest in the boreal forest zone. However, based on studies from hemiboreal and temperate regions (Larrieu et al. 2022; Kõrkjas et al. 2021, 2023; Barone et al. 2025; Brazaitytė et al. 2025), habitat trees contribute to forest biodiversity. Thus, considering their relationship with stand age and size, the diversity and abundance of microhabitats can act as proxies for old-growth structural characteristics, and some, such as fire scars, are indicative of past natural disturbances. We therefore recommend the use of this complementary criterion if regional lists are available of the key microhabitat types known to support biological diversity.

#### 7. Indicator species

Approximately 10% of boreal forest–dwelling multicellular species are confined to OGFs (Hanski 2000). Their association with OGF arises from complex mechanisms, including site history, long-term habitat continuity, dependence on specific microhabitats or host species, and local population dynamics (e.g. Nordén et al. 2014). When such associations are well established, species presence can serve as a robust indicator of OGF attributes. Many of these species in Europe are also regionally red-listed or protected, reinforcing their value for identifying and conserving OGF habitats. Table S2 provides an example of a regional (Estonia) list of OGF indicator species.

For several reasons, no single list of such species can be provided for the entire boreal zone. Species distributions encompass the region to varying extents, and substantial regional differences remain in taxonomic knowledge and recording intensity. Ecologically, the set of species confined to OGF may also depend on the condition of surrounding forests. In intensively managed landscapes, the most demanding species may already be extirpated, and less demanding species may become restricted to the remaining OGF (Lõhmus &Lõhmus 2019). In addition, strong apparent affinities to OGF may emerge in specific regions where old-growth conditions buffer other limiting factors, such as climate, competition, or predation (Lõhmus et al. 2023).

### Implementation

As clearly stated in the EU guidelines, the national implementation should be science-based and not taking economic or political aspects into account. As explained in the theoretical background it is a clear strength that the definition of OGF rests on a combination of mandatory and complementary indicators that *collectively* aim to exclude non-OGFs, and at the same time, do not risk excluding OGFs.

We consider that the current data and knowledge is sufficient to set thresholds conditions required for identification of OGF for the following indicators:

- Native tree species represent a foundation for boreal and hemiboreal biodiversity. Some natural northward migration due to climate change may occur of southern deciduous species, but this does not represent a problem for the definition of OGF as being native. The occurrence of exotic, invasive species may be a potential issue but rather relates to conservation management than representing a problem for identifying OGF.
- Dead wood thresholds based on the proportion of coarse dead wood in relation to total tree volume and taking dead wood heterogeneity into account will better capture OGF conditions than absolute volume thresholds. Established methods to estimate volumes of living and standing dead trees exist as these variables are regularly recorded in forest monitoring and research projects. However, measurement of downed deadwood volume is more challenging and requires harmonization.
- Presence of old trees indicate long term continuity (several tree generations) while the age of a dominant tree cohort risks missing sites subject to recent natural disturbances. Although large trees may be old, care should be taken when estimating number of old trees as also small, supressed trees and slow growing trees can reach high age and indicate stand continuity -a typical feature of especially spruce forests and low productive bog forests. Directly analysing tree ages by means of dendrochronological methods provides the most reliable estimates. However, also more rapid indirect approaches based on morphological traits (e.g., bark texture, branch thickness, crown structure) can also provide reasonable age estimation (Handegard et al. 2021).
- Natural regeneration is clearly dominant in boreal and hemiboreal OGF forests. Although a few examples of old planted stands do occur, most of these have been subject to historical management and only in some rare exceptions may be considered OGF.
- Habitat trees do represent important structures and with numerous species associated with hollow trees, old snags, fire scars etc (Bütler et al. 2024). Given their occurrence on old, large and dead trees they serve as good indicators of OGF conditions. At present there is no relevant data available to identify quantitative thresholds, but their presence provides qualitative information.

For some indicators there is a need for additional consideration before they can be fully implemented in identifying OGF. However, none of these represent a significant obstacle to initiate large scale mapping of OGF.

- Structural complexity is typical for OGFs and clearly linked to natural disturbance processes. An obvious approach to use this indicator is to utilize existing or to develop new field protocols that describe the tree species composition by tree layers, including horizontal variation in tree density. Although examples of this do exist (e.g., Kurlavicius et al. 2004) complete mapping of large areas will be resource demanding. Fortunately, several recent studies using remote sensing have shown that stand structural variability can be estimated by aerial lidar data and support identification of high conservation value forests across large areas (cf. Ørka et al. 2022; Ceccherini et al. 2023). For instance, variation in tree height was a strong predictor for high conservation value forest in the north and south boreal regions of Sweden (Bubnicki et al. 2024). This approach has been found effective even at rather low scan densities (e.g., 2-3 points/m2; Martin and Valeria 2022). Although these methods are readily available and supported by use of relevant statistical estimates, they need further evaluation to ensure that thresholds are set in relation to benchmark OGF areas.
- There is a significant knowledge base on species that are OGF-dependent. However, indicator species suitable for identifying OGFs need to be defined nationally and for different forest types. As all countries in the region have dedicated expert groups for Red-listing species we recommend that they could be tasked to develop lists of OGF indicator species.

While mapping small remnant stands of OGF pose a practical challenge, it should be noted that stand size is not an explicit criterion for the identification of OGF and small areas may provide important refugia for threatened species and act as critical nodes in landscape connectivity (e.g., Wang et al. 2025). When implementing the stand-scale criteria, it is critical to assess how stand and plot size influence quantitative field estimates. Given the typical heterogeneity of OGF, a small stand/plot may not meet threshold values despite representing a remnant OGF. For instance, random small-scale variation in tree mortality may have resulted in lack of certain dead wood types or limited occurrence of old large trees. Such temporal variation may not capture the long historical continuity of small stands/plots. If small remnant patches carry a historical legacy from past conditions, qualitative criteria such as occurrence of habitat trees and indicator species may become particularly important.

## Discussion

Halting biodiversity loss in Europe is a main environmental objective of the EU Biodiversity Strategy to which all Member States are committed. Strictly protecting the remaining primary and old-growth forests is a central action of the strategy, and thus to adopt science-based operational indicators to identify and map them are crucial. If Member States fail to do so they run political, economic, societal, environmental and legal risks. Although the identification of primary and old-growth forests is grounded in ecological science, its implementation is inherently a governance process operating at the interface of science, policy, and land use. In the European Union, old-growth forest indicators function as regulatory instruments that determine legal compliance with biodiversity, climate, and energy legislation, and therefore directly shape forest governance outcomes (Sabatini et□al. 2020; Svensson et□al. 2025). Consequently, operational criteria, threshold setting, and interpretation influence not only conservation effectiveness but also have multiple societal dimensions.

Forest governance in the EU is characterized by divided institutional and often contradictory responsibilities (e.g. Harrinkari et al. 2016; Wolfslerner et al. 2020; Vihma and Toikka 2021). OGF identification sits at this contested boundary, creating incentives for sector-specific interpretations of ecological indicators. Where identification methods lack transparency or scientific robustness, this fragmentation increases the risk of implementation that undermines EU conservation objectives and generates legal and societal conflict (Angelstam et al. 2020; Sabatini et al. 2020).

Forest owners and the forest industry emphasize predictability, legal certainty, and economic continuity, while conservation actors prioritize avoiding biodiversity loss (e.g. Harrinkari et al. 2016). Rural and forest-dependent communities often hold mixed interests, combining economic reliance on forestry with cultural attachments to forest landscapes and long-term ecosystem integrity (Dawson & Angelstam 2026). These perspectives imply asymmetric perceptions of classification errors: false positives are primarily framed as short-term economic costs, whereas false negatives entail long-term ecological, cultural, and social losses. This asymmetry reinforces the need for precautionary, ecologically grounded identification approaches that minimize irreversible outcomes (Wang 2019).

Cultural and traditional knowledge adds an additional dimension to forest governance in Fennoscandia. Indigenous Sámi livelihoods, particularly reindeer herding, depend on forest continuity and structural features commonly associated with old-growth conditions (Norsted et al. 2025). Although such knowledge does not replace ecological indicators, it complements them by highlighting continuity and cultural values in heterogeneous and historically influenced landscapes (Jansson et□al. 2015).

Taken together, these considerations underscore that OGF identification is both a scientific and institutional task. Ecologically robust and transparent operational criteria support effective governance by reducing conflict, improving accountability, and strengthening social legitimacy. Conversely, weak or selectively interpreted criteria risk institutional failure, erosion of trust, and continued biodiversity loss. Integrating ecological evidence with an explicit awareness of governance and societal implications is therefore essential for durable and credible protection of Europe’s remaining OGFs.

A prime example of a weak or selectively interpreted criteria is the Finnish government’s adopted operational criteria for OGFs in Finland, which clearly fail scientific standards, do not meet EU legal requirements, ignore ecological realities, and appear designed to minimize protection.

Consequently, they exclude substantial areas that qualify as OGFs under ther Commission guidelines (Finnish Environment Institute 2024). Major deficiencies include groundlessly high threshold values for the dead wood volume indicator, reliance on mean age of dominating tree cohort and scientifically unfounded minimum area requirements. For example, applying the Finnish Government threshold for dead wood volume in southern boreal pine-dominated forest (minimum 40 m^3^/ha) would have missed >50 % of OGF stands in Siitonen et al. (2026) data and some 45 % in Syrjänen et al. (2024) data. Moreover, the Finnish Government set 4 ha as the minimum area requirement for OGF, which is not based on the Commission guidelines. Together with the ecologically unfounded criterion that dominating tree cohort in OGFs must be older than 140 years, it is highly likely that applying the Finnish Government set of threshold values will miss most pine dominated OGFs in southern boreal forests. Comparable shortcomings occur in Sweden, where new regulations adopt high stand-age (basal area weighted mean age) and area thresholds likely to omit significant OGF areas from strict protection (Swedish Forest Agency 2025; SFS 2025). Failure to safeguard remaining OGFs increases future economic costs e.g. from restoration, harms the forest sector’s reputation, perpetuates conflicts over forest use, accelerates biodiversity loss, and risks EU legal sanctions.

Claims that climate change diminishes the conservation value of OGFs are not supported by current evidence (Watson 2018). Although climate change will alter northern OGFs, their ecological value will persist because long-term structural legacies and native species assemblages enable natural processes to continue under shifting conditions. The value of OGFs is further supported by ensuring species dispersal and colonization of new habitats. Ensuring large-scale connectivity is therefore critical, linking remaining OGFs with production forests (Undin et al. 2024). Continuous cover forestry (Peura et al. 2018) and extended rotations (Roberge et al. 2016) provide practical tools for this integration.

The Commission guidelines distinguish between main and complementary indicators. Main indicators are mandatory, and thus, considered representing characteristics ubiquitous in OGFs. This is logically sound in theory but may cause problems in practice as our assessment above on deadwood thresholds indicates where conditions in OGF and non-OGF overlap. This emphasizes the importance of the complementary indicators in order to avoid false positives and in practice these may be of similar importance as the mandatory indicators.

Indicator-based approaches apply quantitative thresholds to ecosystem characteristics that are inherently continuous and variable. While such thresholds are necessary for operational implementation, they inevitably involve simplifications and somewhat subjective decision boundaries, particularly given limited availability of comprehensive benchmark data on the full variation of old-growth and mature managed forests across regions and site types. Furthermore, as we have theoretically explained - any thresholds inevitably miss some stands that provide OGF functions. For these reasons, ‘old-growthness’ should be understood as a continuum rather than a fixed state, and identification procedures should combine quantitative criteria with qualitative indicators and informed expert judgement. Our threshold choices deliberately err on the side of precaution, reflecting both the EU requirement to protect all remaining OGFs and the recognition that the remaining OGFs are irreplaceable and foundational to long-term ecological, cultural, and societal wellbeing.

