## Supplementary information for "From guidelines to practice: Operational criteria for identifying old-growth forests in northern Europe"

*Ambio*

This supplementary information has not been peer reviewed

Table S1. Volume proportions (%) of native tree species across Fennoscandia and the Baltic states (FAO 2020). * <0.1 %

| **Tree species** | **Total region** | **Norway** | **Sweden** | **Finland** | **Estonia** | **Latvia** | **Lithuania** |
|---|---|---|---|---|---|---|---|
| Scots pine, *Pinus sylvestris* | 39.7 | 29.4 | 37.8 | 50.1 | 30.6 | 33.0 | 36.2 |
| Norway spruce, *Picea abies* | 36.0 | 42.6 | 39.9 | 29.9 | 22.7 | 19.8 | 22.2 |
| Birch, Betula spp. | 16.3 | 17.4 | 15.1 | 16.6 | 25.8 | 23.3 | 15.5 |
| Aspen, *Populus tremula* | 2.6 | 1.8 | 1.5 | 1.7 | 6.7 | 10.1 | 5.8 |
| Black alder, *Alnus glutinosa* | 2.6 | * | 1.1 | 0.3 | 4.5 | 6.5 | 9.4 |
| Grey alder, *Alnus incana* | 2.0 | 1.9 | 1.1 | 0.7 | 6.7 | 5.3 | 3.7 |
| Oak, *Quercus robur* | 0.8 | 0.8 | 1.1 | * | * | 0.4 | 1.0 |
| Willows, *Salix* spp. | 0.5 | 1.1 | 0.4 | 0.3 | 0.6 | * | * |
| Ash, *Fraxinus excelsior* | 0.4 | 0.3 | * | * | 0.8 | 0.7 | * |
| Rowan, *Sorbus aucuparia* | 0.3 | 1.0 | 0.2 | 0.3 | * | * | * |
| Other native species | 0.8 | 3.4 | 1.4 | 0.3 | 1.4 | 0.6 | 0.7 |
| **Total native species** | **100** | **100** | **100** | **100** | **100** | **100** | **100** |

**Table S2.** Volumes ($m^3$/ha) of coarse woody debris (at least 10 cm in diameter, except Siitonen et al. 2026 >5 cm) in different boreal forest vegetation zones of primary old growth forests;. Mean values, standard deviation and 10th percentile (decile) in primary old growth spruce- and pine-dominated (and deciduous dominated hemiboreal) forests in three large compilations of OGFs (n = number of stands; * recalculated from original data).

| Boreal vegetation subzones | n | mean | std | 1st decile | Source |
|---|---|---|---|---|---|
| | Deciduous dominated | | | | |
| Hemiboreal | 35 | 119 | 64 | 48 | Rosenvald et al. 2026* |
| Southern and middle boreal | 22 | 33 | 26 | 5 | Siitonen et al. 2026 |
| | Spruce dominated | | | | |
| Hemiboreal | 50 | 154 | 73 | 66 | Rosenvald et al. 2026* |
| Hemi- and southern boreal | 26 | 138 | 67 | 58 | |
| Middle boreal | 38 | 97 | 44 | 44 | Syrjänen et al. 2024 |
| Northern boreal | 43 | 44 | 25 | 18 | |
| Southern and middle boreal | 121 | 73 | 54 | 12 | Siitonen et al. 2026 |
| | Pine dominated | | | | |
| Hemiboreal | 37 | 78 | 51 | 13 | Rosenvald et al. 2026* |
| Hemi- and southern boreal | 10 | 43 | 25 | 5 | |
| Middle boreal | 18 | 88 | 27 | 49 | Syrjänen et al 2024 |
| Northern boreal | 22 | 21 | 15 | 7 | |
| Southern and middle boreal | 90 | 37 | 29 | 7 | Siitonen et al. 2026 |

**Table S2.** Example of a regional evidence-based list of indicator species of old-growth forests: Estonia.

| Species | Group | Red-list status | Evidence of old-growth specificity | References |
|---|---|---|---|---|
| *Amylocystis lapponica* | Wood-inhabiting basidiomycetes | CR | Distributed only in, and spreading along, old-growth reserve networks | [1,2,3] |
| *Antrodia piceata* | | EN | Only recorded in old natural forests where occurs regularly | [1,3] |
| *Fomitopsis rosea* | | NT | Substrate- and connectivity-dependent species on coarse spruce wood | [4] |
| *Phellinus nigrolimitatus* | | LC | Substrate-, stand-age and connectivity-dependent species on coarse spruce wood | [4] |
| *Rigidoporus crocatus* | | LC | Confined to fallen trunks in permanently wet conditions, mostly in natural swamp forests | [1,3] |
| *Phellinus chrysoloma* | Tree-parasitic basidiomycetes | LC | Microhabitat-creating species on slowly grown old live spruce | [3,4] |
| *Phellinus pini* | | LC | Hollow-creating keystone species in old live pine trees | [3,4,5,7] |
| *Junghuhnia pseudozilingiana* | Saprotrophic basidiomycete | VU | On old aspens hollowed by *Phellinus tremulae*, only in closed-canopy forests | [6,7] |
| *Hydnellum aurantiacum* | Ectomycorrhizal basidiomycete | VU | Disturbance-sensitive soil fungus in old conifer forests | [7] |
| *Multiclavula mucida* (basidiocarps) | Wood-inhabiting basidiolichen | VU | On large moist decayed fallen trunks in naturally developed forests | [7,9] |
| *Alectoria sarmentosa* | Pendulous canopy-dwelling macrolichen | VU | Confined to natural mixed and conifer forests with high temporal connectivity | [1] |
| *Lobaria pulmonaria* | Epiphytic macrolichens on live deciduous trees | VU | Connectivity-dependent species on nemoral broad-leaved trees and aspen; typically in forests >100 years old | [7,8,9] |
| *Menegazzia terebrata* | | VU | Confined to hydrologically intact natural wet forests (apparently connectivity-dependent) | [7,9] |
| *Nephroma laevigatum; N. bellum* | | EN | Confined to old deciduous trees in natural closed-canopy forests | [1,10] |
| *Mycoblastus sanguinarius* | Bark- and wood-inhabiting lichens | LC | Confined to old moist forests | [9] |
| *Xylopsora friesii* | | NT | In conifer forests with continuity of small-scale (fire) disturbances and slow-grown old trees | [9,11] |
| *Chaenotheca gracilenta* | Microhabitat-dependent microlichen | EN | Confined to shady moist microhabitats on dying and uprooted old trees in natural forests | [7,9,12] |
| *Buxbaumia viridis* (sporophyte) | Wood-inhabiting bryophyte | VU | Confined to >80 years-old conifer forests with high temporal connectivity | [7,13] |
| *Sphagnum wulfianum* | Ground-dwelling bryophytes | LC | In old spruce forests near mires; competition-sensitive | [1,14] |
| *Bazzabia trilobata* | | NT | In old mixed forests | [1] |

| | | | | |
|---|---|---|---|---|
| *Cucujus cinnaberinus* | Wood-inhabiting beetle | EN | In Estonia recorded exclusively in old-growth stands on dying and dead aspens | [15] |
| *Vertigo alpestris* | Snail | LC | Confined to old stands with abundant coarse woody debris | [16] |
| *Pteromys volans* (occupied cavity) | Cavity-nesting mammal | CR | In Estonia, confined to cavity-rich vertically diverse forests of ca. 120 years optimal age | [17,18] |
